# Systemic Delivery of Human Mesangioblasts mediated by a Nanofiber Scaffold restores Dystrophin Expression in Immunodeficient Dystrophic Mice

**DOI:** 10.64898/2026.03.31.715524

**Authors:** S. Amer, L. Bragg, S. Santoleri, G. Cossu, F Galli

## Abstract

Delivery of cells or vectors in advanced therapies is probably the major challenge for genetic disorders that affect a large part of the body such as Duchenne Muscular Dystrophy (DMD). Here, we describe a novel approach for systemic cell delivery based upon an implantable bio-scaffold composed of aligned polycaprolactone nanofibers coated with laminin, able to support adhesion and extensive proliferation of mesoderm cells both in *vitro* and when implanted subcutaneously in a DMD mouse model. The scaffold is rapidly vascularised leading to cell entering the circulation and colonising multiple distal organs, including distant skeletal muscles and heart. Cells survive in colonized muscles and differentiate into muscle fibres that produce well detectable levels of dystrophin and α-sarcoglycan. These results are game changing for cell therapy, as they allow colonization of life essential but “difficult to reach” muscles such as diaphragm and heart while avoiding invasive catheterization. Once optimised, this approach will rapidly enter clinical experimentation for DMD, other muscular dystrophies, and possibly other genetic disorders of the mesoderm.

Graphical abstract
Study design and therapeutic outcome.
Muscle biopsies were obtained from Duchenne muscular dystrophy (DMD) patients to isolate human DMD mesangioblasts (DMD-hMabs). Cells were genetically corrected using a lentivirus carrying a snRNA able to induce exon skipping (U7snRNA), generating U7-hMabs (1). U7-hMabs were seeded onto laminin-coated polycaprolactone (Lam-PCL) nanofiber scaffolds and implanted into the back muscle of DMD-NSG mice. This platform enabled systemic distribution of hMabs cells through circulation, resulting in engraftment across multiple muscle groups, including tibialis anterior, triceps, diaphragm and heart.

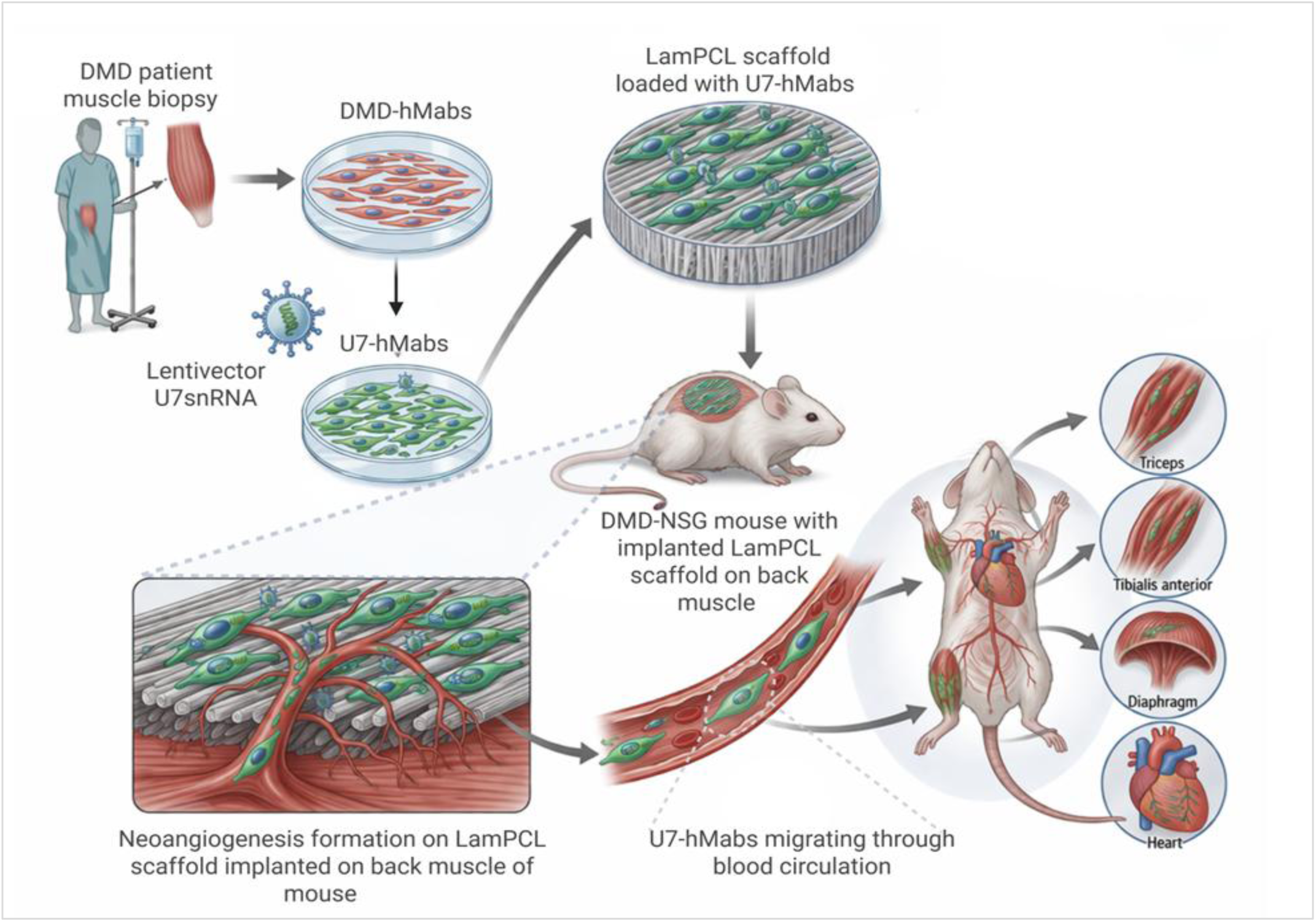

## Introduction

Four decades after the cloning of the dystrophin gene (2), Duchenne Muscular Dystrophy (DMD) remains without an effective therapy. Novel generation drugs aimed at correcting the dystrophin transcript, including Ataluren and exon-skipping oligonucleotides (e.g. Exondys), have yielded limited clinical benefit (3), leading to the withdrawal of marketing authorization for Ataluren (https://www.ema.europa.eu/en/news/translarna-ema-re-confirms-non-renewal-authorisation-duchenne-muscular-dystrophy-medicine).

Adeno-Associated Virus (AAV)–based in *vivo* gene therapies delivering either micro-dystrophin constructs or CRISPR–Cas9 components initially showed promise but have encountered substantial translational barriers. These include immunogenicity, which prevents repeat dosing, pre-existing immunity, limited cargo capacity, high manufacturing costs, and significant toxicity, which caused many serious adverse events, several of which lethal (4–6).

Cell-based therapies, while not associated with comparable systemic toxicity, have similarly failed to demonstrate efficacy in muscular dystrophies, primarily due to inefficient delivery and poor engraftment (7). Following more than fifteen years of preclinical studies in small and large animal models (8–11), we conducted a first-in-human Phase I/IIa clinical trial of cell therapy for DMD (EudraCT 2011-000176-33). Although the trial confirmed safety, therapeutic benefit was minimal (12), largely reflecting extremely low donor-cell engraftment (<1%).

To overcome this limitation, we developed and validated a novel strategy that integrates cell therapy with exon-skipping, whose efficacy was demonstrated in rodent models (1) and subsequently in a proof-of-concept Phase I clinical trial (EudraCT 2023-000148-47) (Cossu et al. in preparation). In this approach, genetically corrected autologous cells act as “Trojan horses”: once incorporated into regenerating myofibres, they express a small nuclear RNA (snRNA) that induces skipping of dystrophin exon 51 not only in donor nuclei but also in neighbouring dystrophic resident nuclei. This amplifying mechanism restores dystrophin expression to therapeutically relevant levels, resulting in complete functional rescue (1).

Despite this advance, effective systemic delivery remains a major challenge in DMD, a disease affecting most skeletal muscles. We previously developed intra-arterial delivery via femoral artery catheterization in mice, dogs, and patients (8, 9, 12). This strategy was optimized for mesangioblasts (Mabs), which, under inflammatory conditions, interact with endothelial adhesion molecules similarly to leukocytes (13). However, whole-body catheterization is invasive, requires general anaesthesia, and poses clinical risks, particularly in the context of repeated administrations (14). Most importantly, essential muscle groups, including diaphragm, heart, and dorsal/postural muscles, are poorly accessed by standard catheterization routes, resulting in very little engraftment.

Here, we show that a subcutaneously implanted bio-scaffold, consisting of polycaprolactone (PCL) nanofibers, coated with laminin (thereafter referred as “LamPCL scaffold”) and seeded with human mesangioblasts (hMabs), becomes rapidly vascularized and functions as a systemic cell-delivery platform. Cells are progressively released into the circulation, enabling widespread body distribution. Upon homing to skeletal muscles, the delivered hMabs differentiate into new myofibres and restore, at least partially, dystrophin and associated protein expression, thereby overcoming key limitations of current delivery strategies.

## Results

### Scaffold characterization and biocompatibility in *vitro*

A polycaprolactone (PCL) scaffold, fabricated from an FDA-approved synthetic polymer, was developed for its ability to mimic the elasticity of the native extracellular matrix (ECM) of skeletal muscle (15, 16). Using optimized electrospinning parameters, we fabricated an aligned nanofiber scaffold displaying a smooth and uniform surface, as showed in SEM micrographs (Figure 1a). Fibre diameters averaged 723 ± 342 nm (Figure 1b), confirming overall uniformity of the nanofiber architecture. To enhance biocompatibility and cellular adhesion, the PCL scaffold was coated with laminin (5 ug/ml in PBS) (LamPCL scaffold).

**Figure. 1:**
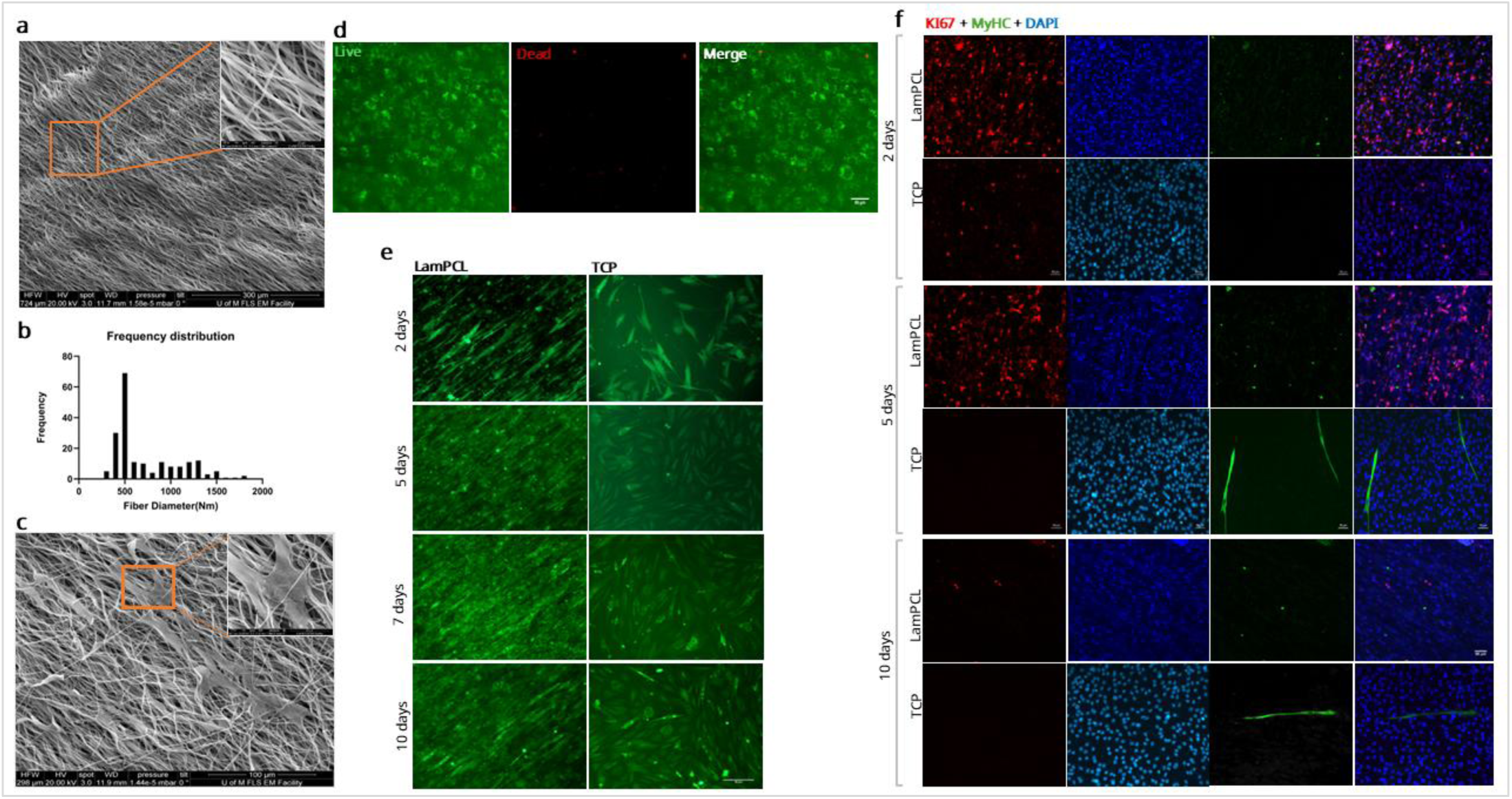
Scaffold characterization, biocompatibility, and human mesangioblasts proliferation rate in *vitro.* (**a**). Representative scanning electron microscope micrographs of aligned LamPCL nanofibers. Scale bars= 300µm, (magnification on the upper right corner = 20µm). (**b**). **Histogram** illustrating the distribution of fibre diameters in electro-spun aligned LamPCL scaffold, n= 3 (electrospun fibres sheet); 60 fibres for each sample. (**c**) **Cell morphology**, representative scanning electron microscope micrographs of cultured wild type human mesangioblasts after 24 hours on LamPCL nanofiber, showing the distribution and flatten morphology of cells over the scaffold, scale bar=100 µm, (magnification on the upper right corner, scale bar= 20 µm). (**d**) **Cell viability assay**, representative confocal images of Live (green)- Dead (red) staining test of Wt-hMabs on electro-spun LamPCL aligned nanofiber, n=3, scale bar =50µm. (**e**). **Cell morphology, distribution and alignment on scaffold**, representative live microscopic images of GFP^+^ Wt-hMabs seeded on LamPCL nanofiber scaffold over 2 days, 5 days, 7 days, and 10 days after cell seeding, scale bar =50µm. (**f**). **Wt-hMabs proliferation/differentiation assay**, representative confocal microscopy images of cultured Wt-hMabs in growth medium, stained with anti-Ki67 (red), anti-Myosin heavy chain (MyHC) (green) antibodies, and DAPI (blue), at 2 days, 5 days, and 10 days. LamPCL (Laminin-PCL scaffold), TCP (Tissue culture plate_ control), scale bar =50µm.

Scanning electron microscopy was performed 24 hours after the plating Wilde type hMabs (Wt-hMabs) on the LamPCL scaffold to evaluate cell adhesion. SEM micrographs confirm attachment of cells on the scaffold (Figure 1c).

Cell viability was assessed using Live/Dead staining 24 hours after seeding. Wt-hMabs exhibit widespread distribution across the scaffold and predominantly green positive cells indicating high viability. Only a small portion of dead cells, stained in red, was detected corresponding to an average viability rate of approximately 97±0.8% (Figure 1d). To examine cell organization over time, GFP positive (GFP^+^) Wt-hMabs were seeded on the LamPCL scaffold and analysed at 24 hours, 2 days, and 10 days. At all-time points, cells remained uniformly distributed and oriented in line with the nanofiber’s alignment. Notably, even after 10 days, the GFP^+^ Wt-hMabs maintained their alignment, demonstrating integration and long-term survival within the LamPCL scaffold (Figure 1e).

Proliferative capacity was evaluated by immunofluorescence (IF) for Ki67 in Wt-hMabs cultured on LamPCL scaffold and maintained in growth medium (Figure 1f). Cells were analysed at days 2, 5 and 10. Quantification, by counting Ki67-positive (Ki67^+^) cells, showed active proliferation at Day 2, reached a peak of proliferation at Days 5, and showed a marked decline by Day 10 (Supp. Figure 1). These results suggest an early phase of active expansion on the scaffold followed by a transition to reduced cycling, not substantially different from what happens in tissue culture dishes.

### In *vivo* scaffold biocompatibility and integration with host muscle

After *in vitro* analysis, we implanted subcutaneously (SC) the LamPCL scaffolds to evaluate their ability to support Wt-hMabs survival, proliferation, and long-term integration within adjacent (back) muscle. Four weeks after SC implantation of the LamPCL scaffold in Wild type immunodeficient NSG mice (Wt-NSG), the scaffold and the back muscle were explanted for analysis. IF for Ki67 showed high cell proliferation within the scaffold, while Myosin Heavy Chain (MyHC) staining showed absence of myogenic differentiation at this stage (Figure 2a, 2b). Importantly, IF for human Lamin AC confirmed persistence of Wt-hMabs within the scaffold (Figure 2c) that had not be replaced by mouse cells. Together, these findings indicate that the LamPCL scaffold supports long-term survival and proliferation of transplanted cells, *in vivo*, without inducing premature *in situ* differentiation, thereby preserving their migratory potential.

**Figure. 2:**
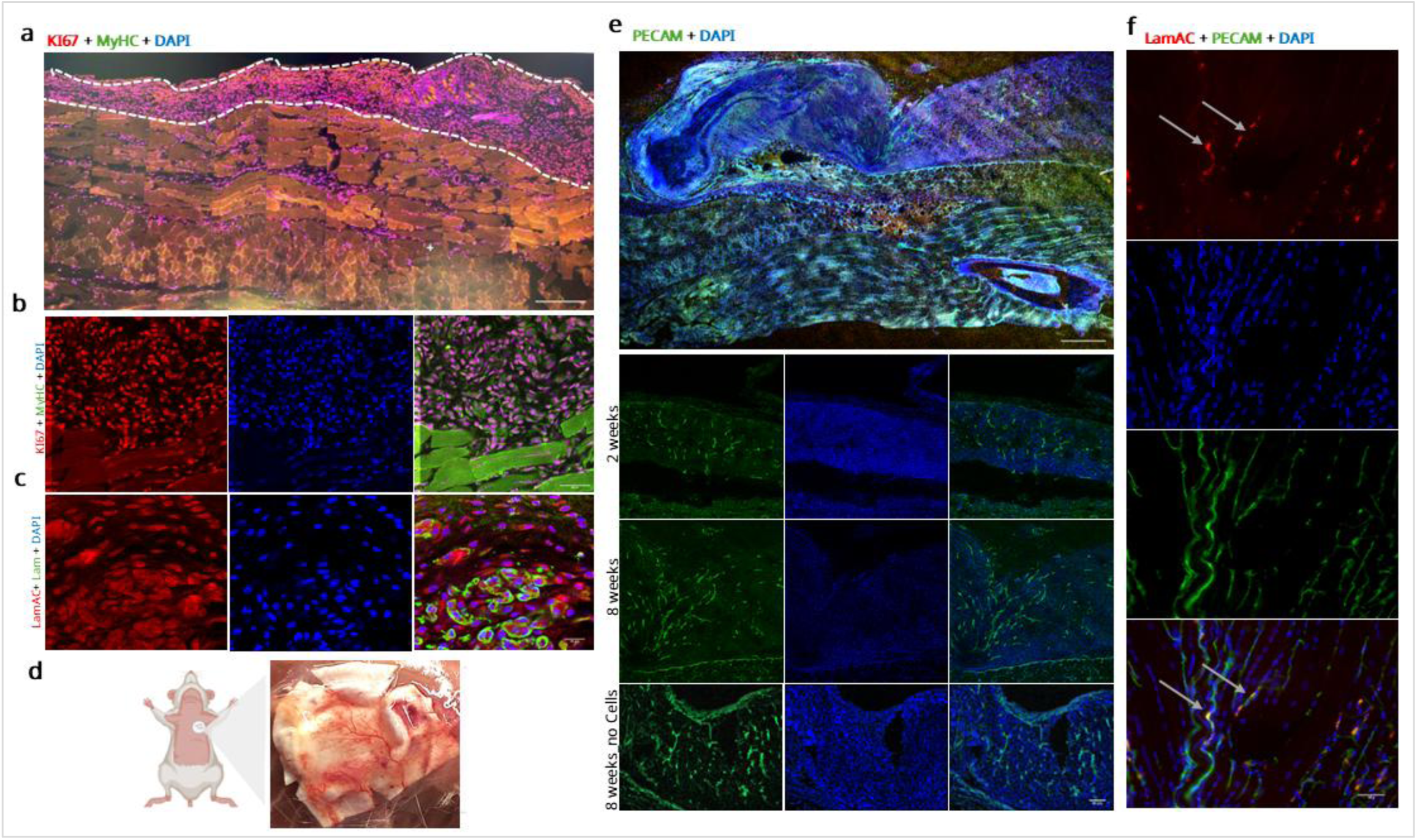
In *vivo* scaffold biocompatibility and integration with muscles of Wt-NSG mice. (**a**). Representative overview confocal microscopy images of a cross-sectioned back muscle with the implanted scaffold loaded with wild type human mesangioblasts positioned on top stained with anti-Ki67 (red), anti-Myosin heavy chain (MyHC) (green) antibodies, and DAPI (blue). The scaffold area is outlined with white dashed lines, scale bar = 50µm. (**b**). Higher magnification of scaffold area on top of back muscle stained with anti-Ki67 (red), anti-Myosin heavy chain (MyHC) (green) antibodies, and DAPI (blue), scale bar=50µm. (**c**). Higher magnification of scaffold area on top of back muscle stained with anti-Lamin AC (red), anti-Laminin (green) antibodies and DAPI (blue), scale bar=20µm. (**d**). Graphical illustration showing the implanted scaffold over the mouse back muscle; gross examination showed blood vessels growth into the scaffold. (**e**). Representative confocal overview images of a cross-sectioned back muscle with the implanted scaffold loaded with Wt-hMabs positioned on top, showing neovascularization of the scaffold area stained with anti-PECAM (green) antibody, and DAPI (blue) 8 weeks after transplantation, scale bar = 50µm. (**e**). Representative confocal microscopy images of a cross-sectioned back muscle with the implanted scaffold loaded Wt-hMabs and stained with anti-PECAM (green) antibody, and DAPI (blue) after 2 and 8 weeks; a cell free scaffold (is also shown at 8 week) transplantation. Scale bar = 50µm. (**f**). Representative confocal images of Wt-hMabs stained with anti-Lamin AC (Red) antibody within the wall of blood vessel stained with anti-PECAM (Green) antibody and DAPI (Blue) in longitudinal section of the tibialis anterior muscle, scale bar = 50µm. **Laminin = Lam, Lamin AC = LamAC.**

Interesting, macroscopic examination of the implant site revealed the presence of abundant newly formed blood vessel within the LamPCL scaffold (Figure 2d). To evaluate scaffold-driven angiogenesis, we performed IF analysis using PECAM (CD31), a marker of endothelial cells and neovascularization. Two weeks after implantation, we detected PECAM positive cells within the scaffold, indicating the occurrence of angiogenesis. After four weeks, PECAM staining increased in both density and structural organization, confirming robust angiogenesis of the LamPCL scaffold (Figure 2e).

Importantly, in the control group implanted with LamPCL scaffold devoid of Wt-hMabs, comparable PECAM-positive vascular structures were observed, suggesting that neovascularization is induced by the LamPCL scaffold independently from Wt-hMabs released signals (Figure 2e). These results indicate effective scaffold integration into host tissue and highlight its ability to promote angiogenesis, that likely supports cell survival and proliferation.

Finally, confocal imaging of longitudinal sections of *Tibialis Anterior* (TA) muscle revealed human Lamin AC positive (Lamin AC^+^) nuclei, corresponding Wt-hMabs, associated with PECAM positive vascular structures (Figure 2f), providing direct evidence that transplanted Wt-hMabs enter the circulation thus enabling body-wide distribution from a single implantation site; indeed we detected human cells inside the blood vessels of all muscles we have looked at, as well as in filter organs (detailed below).

### Wt-hMabs migration after induced muscle injury

Migration of cells from scaffold implantation was first evaluated in a WT-NSG mouse model in the absence of induced muscle injury to establish a baseline condition. Histological analysis of the underlying back muscle revealed absence of human Lamin AC^+^ cells after 4 weeks of implantation indicating that Wt-hMabs remained confined to the blood vessels and did not migrate into intact host tissues (Supp. Figure 2), supporting the notion that, like leukocytes, they need inflammation to extravasate (8).

A more detailed histological examination, however, identified localized mechanical damage induced during scaffold fixation by the suture needle. Histological analysis of cross-sections revealed focal areas of muscle damage directly beneath the implanted scaffold (Figure 3a). Higher-magnification confocal imaging revealed Wt-hMabs, human Lamin AC^+^, centrally nucleated within newly formed, regenerating myofibres (Figure 3b), indicating that Wt-hMabs had migrated from the scaffold into the damaged area and contributed to muscle regeneration. To validate this observation under more controlled and reproducible conditions, a defined injury model was subsequently employed. Wt-hMabs-seeded LamPCL scaffolds were implanted 24h after cardiotoxin (CTX) injection into the underlying back muscle. Confocal imaging revealed a clear anatomical separation between the LamPCL scaffold and host muscle (dashed line in Figure 3c), with Wt-hMabs distributed both within the scaffold and in the adjacent regenerating muscle tissue.

**Figure. 3:**
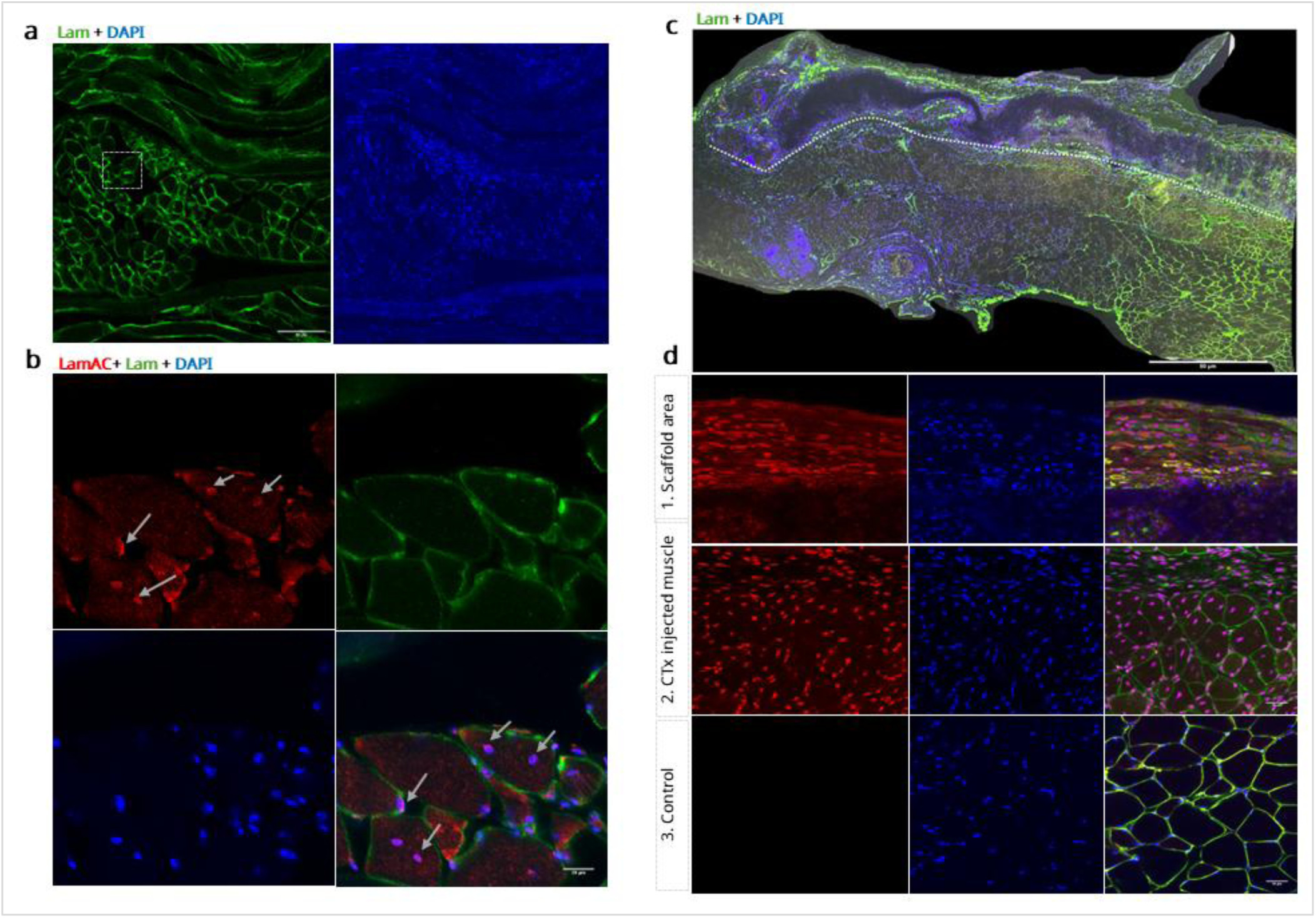
Cell migration after induced muscle injury. (**a**). Representative confocal microscopy images of a cross-sectioned back muscle with the implanted scaffold loaded with wild type human mesangioblasts (Wt-hMabs) in Wt-NSG mouse, showing localized damaged area (white dashed square), stained with anti-Laminin (green) antibody, and DAPI (blue) 4 weeks after transplantation, scale bar = 50µm. (**b**). Representative confocal microscopy images of higher magnification of area with localized damage showing newly regenerated myofibres with centrally located human nuclei (white arrows) stained with anti-Lamin AC (Red), anti-Laminin (green) antibodies and DAPI (blue) 4 weeks after transplantation, scale bar = 20µm. (**c**). Representative overview confocal microscopy image of cross-sectioned back muscle in Wt-NSG mouse with acute injury induced by cardiotoxin injection (CTX), 4 weeks after transplantation of a scaffold loaded with Wt-hMabs; the figure shows wide distribution of human cells across the CTX injected muscle, stained with anti-Lamin AC (red), anti-Laminin (green) antibodies and DAPI (blue), scale bar = 50µm. (**d**). Representative high magnification confocal microscopy images of a cross-sectioned implanted scaffold loaded with Wt-hMabs, positioned on top of the back muscle in Wt-NSG mouse with CTX injury. 1.Top part of the scaffold, showing presence of human cells stained with anti-Lamin AC (red) aligned with nanofiber structure, 2. Regenerated damaged muscle by CTX showing localization of centrally nucleated Lamin AC-positive myofibres. 3. Muscle sections of control Wt-NSG mouse with neither scaffold implantation nor CTX injection showing absence of human cells localization in mouse muscles, Lamin AC (red), Laminin (green), DAPI (blue), scale bar = 50µm. **Laminin = Lam, Lamin AC = LamAC.**

Further analysis of the scaffold revealed Wt-hMabs aligned along the nanofiber architecture, consistent with scaffold-guided cell organization (Figure 3d). In the underlying CTX-injured muscle, numerous centrally nucleated myofibres expressing human Lamin AC, corresponding to Wt-hMabs, were observed, together with significant variability in fibre diameter (Figure 3d). These hallmarks of active regeneration indicate that Wt-hMabs migrated from the LamPCL scaffold and fused with regenerating host myofibres. In contrast, muscle from control group that did not receive CTX-induced injury displayed healthy, uniformly sized myofibres and no detectable human Lamin AC^+^ cells (Figure 3d). Collectively, the absence of donor Wt-hMabs in uninjured muscle confirms that cell migration from the LamPCL scaffold is injury-dependent, rather than the results of spontaneous distribution in every tissue.

### hMabs migration and dystrophin restoration in dystrophic mice

Given the intrinsic ability of Mabs to transmigrate across blood vessel walls in the presence of inflammation (12, 17), together with the results observed after CTX injection, we investigated whether Wt-hMabs delivered via the LamPCL scaffold could migrate beyond the implantation site in dystrophic immune deficient mice (DMD-NGS:12). IF revealed widespread distribution of Wt-hMabs throughout the back muscle, where donor cells were associated with clusters of dystrophin-expressing fibres, indicating successful engraftment and differentiation (Figure 4a).

**Figure. 4:**
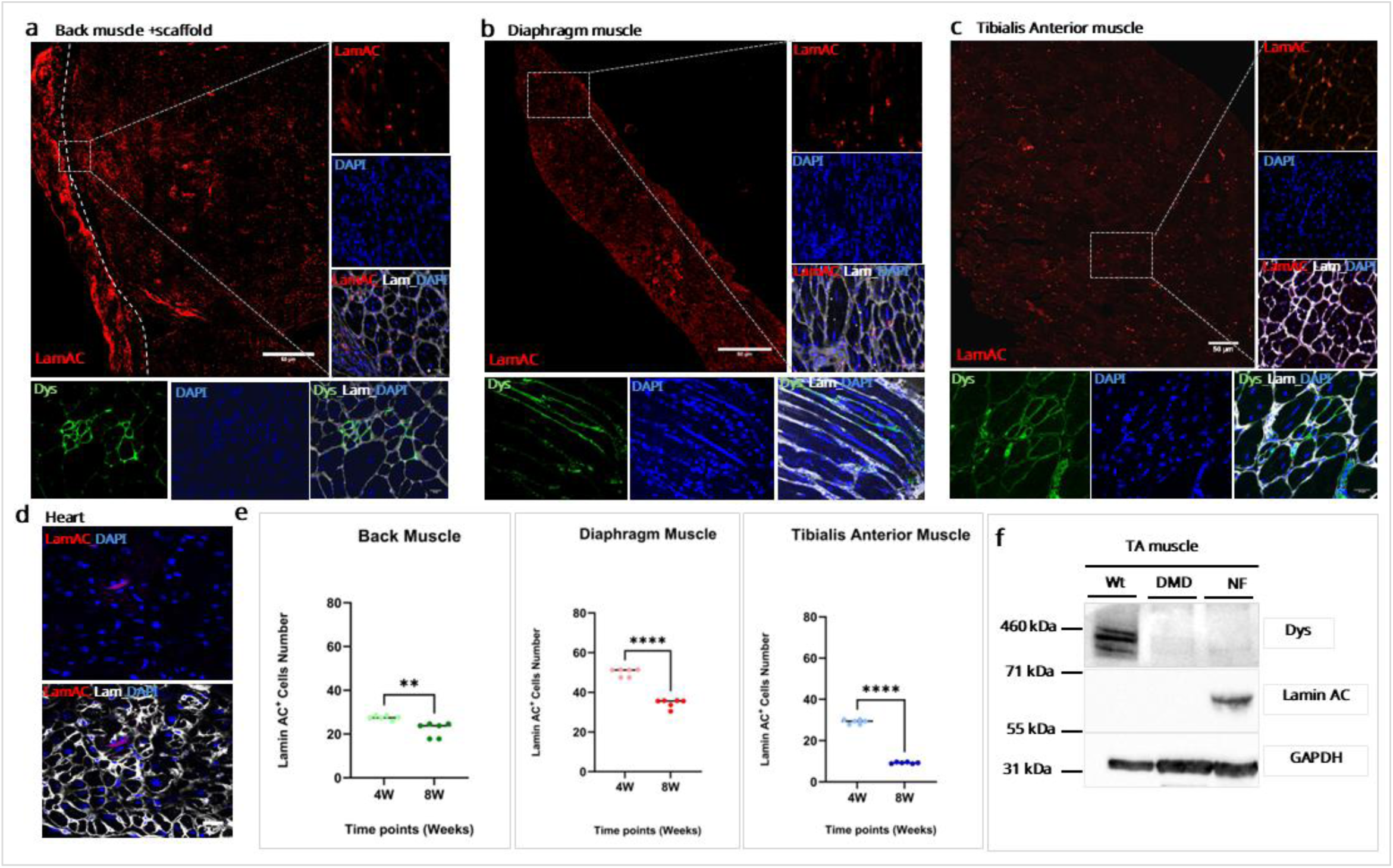
Engraftment and dystrophin restoration four weeks after transplantation of wild type human mesangioblast–loaded (Wt-hMabs) scaffolds in DMD-NSG. (**a**). Representative overview confocal microscopy images of a cross-sectioned back muscle, showing distribution transplanted Wt-hMabs of back muscle-stained with anti-Lamin AC (red) antibody, Scale bar = 50µm. *Vertical column*, Higher magnification confocal images of back muscle showing Lamin AC (red), laminin (grey), and DAPI (blue) expressing human cells. *Horizontal row:* confocal images of back muscle of dystrophin (green), Laminin (grey) and DAPI (blue), scale bar = 20µm. (**b**). Representative overview confocal microscopy images of a cross-sectioned diaphragm muscle, showing distribution of transplanted Wt-hMabs, stained with anti-Lamin AC (red) antibody, scale bar = 50µm. *Vertical column,* Higher magnification confocal images of diaphragm muscle stained with anti-Lamin AC (red), anti-laminin (grey), and DAPI (blue). *Horizontal row:* confocal images of diaphragm muscle of dystrophin (green), laminin (grey), and DAPI (blue), scale bar = 20µm. (**c**). Representative overview confocal microscopy images of a cross-sectioned Tibialis Anterior (TA) muscle, showing distribution of transplanted Wt-hMabs, scale bar = 50µm. *Vertical column,* Higher magnification confocal images of TA muscle stained with anti-Lamin AC (red), anti-laminin (grey) antibodies and DAPI (blue). *Horizontal row:* confocal images of TA muscle stained with anti-dystrophin (green), anti-laminin (grey), and DAPI (blue), scale bar = 20µm. (**d**). Representative confocal microscopy images of cross-sectioned the Heart, 4 weeks after implantation of Wt-hMabs - loaded scaffold, stained with anti-Lamin AC (Red), anti-Laminin (grey) antibodies and DAPI (Blue), scale bar = 20µm. (**e**). Scatter blot graphs showing the number of Lamin AC^+^ cells distribution, in Back muscle, Diaphragm, Tibialis Anterior at 4-, and 8 weeks after implantation of scaffolds loaded with Wt-hMabs. The y-axis represents the number of Lamin AC^+^ cells/0.2mm^2^, while the x-axis indicates different time points (weeks). Each bar indicates the mean number of cells, n=6 (5 sections each animal) represented by circles. Data analysis was performed using unpaired t-test with Welch’s correction. Statistical significance is denoted by asterisks, with increasing numbers of asterisks indicating higher levels of significance (p < 0.05), error bars = SD; data are presented as mean ± SD (f). Western blot analysis of Lamin AC protein expression in Tibialis Anterior 4 weeks post implantation. **Laminin = Lam, Lamin AC = LamAC, DYS = dystrophin.**

Notably, Wt-hMabs and dystrophin restoration were also detected in the contralateral back muscle (Supp. Figure 3a), suggesting long-range migration from the implantation site. Unexpectedly, extensive colonization of the diaphragm, a muscle typically inaccessible using standard cell distribution approaches, was also observed, with clear presence of Wt-hMabs and robust dystrophin expression (Figure 4b). Similar findings in TA, *Gastrocnemius* (G), and *Triceps* (Tri) muscle (Figure 4c, Supp. Figure 3b, d) further confirmed that Wt-hMabs can be systematically distributed via the newly formed vascular network within the LamPCL scaffold. Importantly, analysis of filter organs, including the liver and kidneys, revealed only few Wt-hMabs (Supp. Figure 3c) indicating that donor cells did not accumulate preferentially in filter organs.

Given the clinical relevance of cardiac involvement in DMD, we also examined the heart. Although Wt-hMabs were detected into cardiac tissue (Figure 4g), the cells did not differentiate into cardiomyocytes, and no dystrophin expression was observed in the heart. This is consistent with a previous report showing that Mabs do not spontaneously undergo cardiomyogenic differentiation (18), indicating that additionally strategies will be required to achieve efficient cardiac targeting.

To determine whether Wt-hMabs engraftment was maintained or modified overtime following scaffold implantation, human Lamin AC was used as a quantitative marker of donor cells. IF showed high numbers of human Lamin AC^+^ cells at 4 weeks post-implantation, followed by a reduction at 8 weeks (Supp. Figure 4a-e). In the back muscle, diaphragm, TA, Tri, and G engraftment at 4 weeks averaged ∼ 27, 50, 29, 60, and 46 cells/ 0.2mm^2^, respectively followed by a significant decline at 8 weeks ∼ 22, 33, 9, 28, and 22 cells/ 0.2mm^2^ (Figure 4c, and Supp Figure 4f). Based on this temporal profile and considering that four weeks post-engraftment a well-established time point for donor cells differentiation and functional protein expression, Western Blot (WB) analysis was performed at four weeks to assess the presence of Wt-hMabs derived dystrophin in TA. WB confirmed the presence of human donor cells exclusively in DMD-NSG mice receiving the LamPCL scaffold, validating the specificity of the delivery protocol (Figure 4f). However, despite detectable donor cells engraftment, dystrophin levels remained below the threshold detectable by WB indicating that the level of engraftment was insufficient to achieve significant dystrophin restoration (Figure 4f).

### U7-hMabs cell migration and dystrophin restoration in dystrophic mice

To overcome the limited dystrophin expression observed with Wt-hMabs, we tested dystrophic hMabs genetically corrected with a lentivector expressing U7snRNA, able to induce skipping of human exon 51 (U7-hMabs). This U7snRNA can diffuse along the myofibre correcting also neighbouring nuclei and thus amplifying dystrophin production well above the level produced by Wt-hMabs (1). U7-hMabs were seeded onto the LamPCL scaffold to evaluate their in *vivo* distribution and therapeutic potential.

Four weeks post-implantation, IF revealed widespread distribution of U7-hMabs (human Lamin AC^+^) throughout the host back muscle, accompanied by numerous dystrophin-positive fibres (Figure 5a). Importantly, U7-hMabs were also detected along the entire diaphragm, where they integrated into muscle fibres and induce dystrophin expression (Figure 5b). Analysis of the TA further confirmed long-range cell distribution, with abundant human Lamin AC^+^ cells indicating high engraftment of U7-hMabs and dystrophin expression throughout the muscle (Figure 5c). Comparable results were also observed in Tri and G (Supp Figure 5a, b). Similar patterns were maintained at 8 weeks post-implantation across multiple muscle groups, indicating sustained engraftment (Supp Figure 6).

**Figure. 5:**
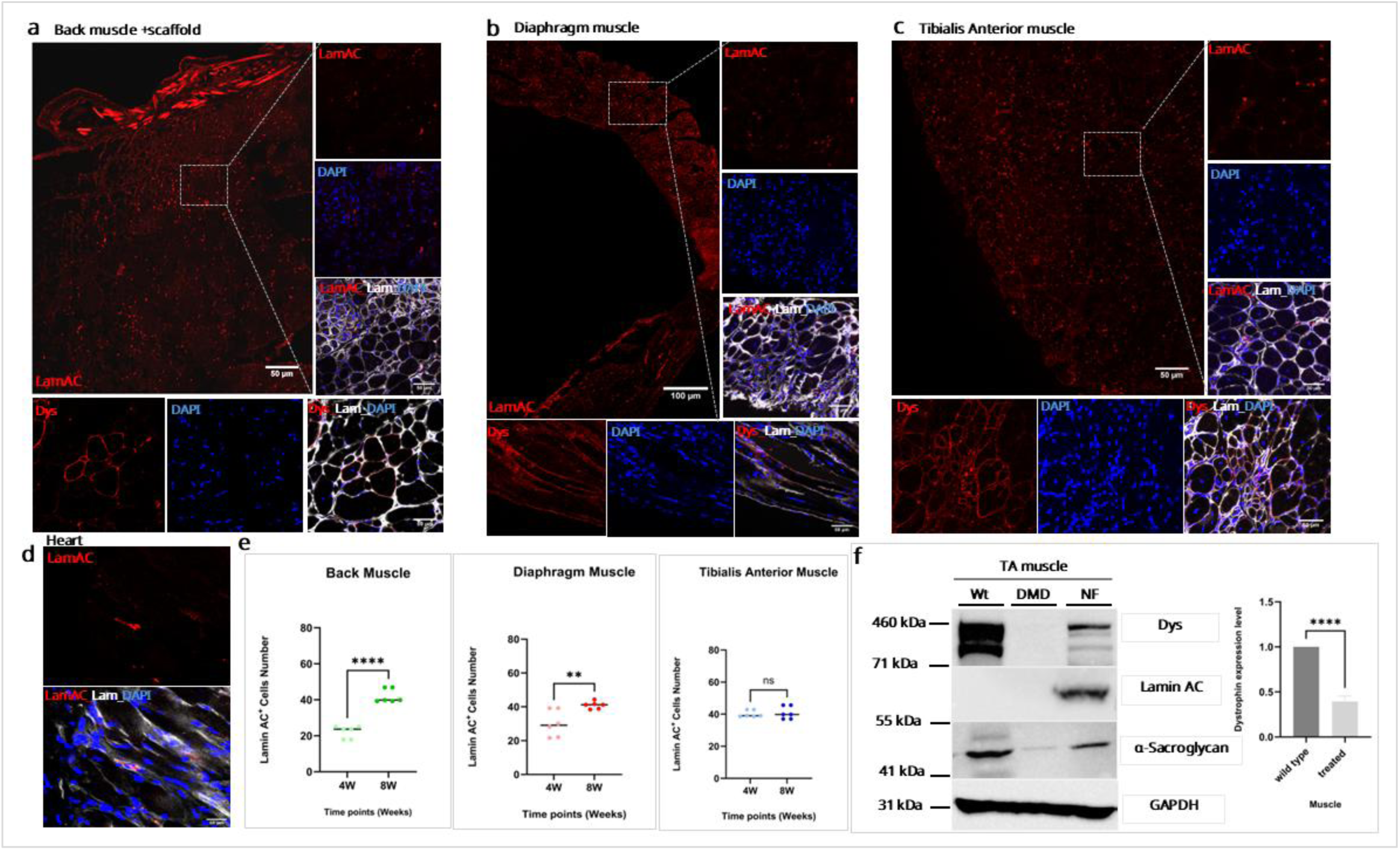
Engraftment and dystrophin restoration four weeks after transplantation scaffolds loaded with dystrophic, U7-genetically corrected human mesangioblasts (U7-hMabs) in DMD-NSG mice. (**a**). Representative overview confocal microscopy images of a cross-sectioned back muscle, showing distribution of transplanted U7-hMabs stained with anti-Lamin AC (red) antibody within the dystrophic muscle 4 weeks after transplantation, scale bar = 50µm. *Vertical column,* Higher magnification confocal images of back muscle stained with anti-Lamin AC (red), anti-laminin (grey) antibodies and DAPI (blue). *Horizontal row:* confocal images of back muscle of stained with anti-dystrophin (red), anti-laminin (grey), and DAPI (blue), scale bar = 20µm. (**b**). Representative overview microscopy images of a cross-sectioned diaphragm muscle, showing distribution of transplanted U7-hMabs stained with anti-Lamin AC (red) antibody; scale bar = 50µm. *Vertical column,* Higher magnification confocal images of diaphragm muscle stained with anti-Lamin AC (red), anti-laminin (grey) antibodies and DAPI (blue). *Horizontal row:* confocal images of diaphragm muscle stained with anti-dystrophin (red), anti-laminin (grey) antibodies, and DAPI (blue), scale bar = 20µm. (**c**). Representative overview confocal microscopy images of a cross-sectioned Tibialis Anterior (TA) muscle, showing distribution of transplanted U7-hMabs stained with anti-Lamin AC (red) antibody; scale bar = 50µm. *Vertical column,* Higher magnification confocal images of TA muscle stained with anti-Lamin AC (red), anti-laminin (grey), and DAPI (blue). *Horizontal row;* confocal images of TA muscle stained with anti-dystrophin (red), anti-laminin (grey) antibodies, and DAPI (blue); scale bar = 20µm. (**d**). Representative confocal microscopy images of cross-sectioned heart, 4 weeks after implantation of U7-Mabs-loaded scaffold. Section was stained with anti-Lamin AC (Red), anti-Laminin (grey), DAPI (Blue), scale bar = 20µm. (**e**). Scatter blot graphs showing the number of Lamin AC^+^ cells distribution, in Back muscle, Diaphragm, Tibialis Anterior at 4-, and 8weeks post-implantation with U7-hMabs cells. The y-axis represents the number of Lamin AC^+^ cells/0.2mm^2^, while the x-axis indicates different time points (weeks). Each bar indicates the mean number of cells, n=6 (5 sections each animal) represented by circles. Data analysis was performed using unpaired t-test with Welch’s correction. Statistical significance is denoted by asterisks, with increasing numbers of asterisks indicating higher levels of significance (p < 0.05), error bars = SD; data are presented as mean ± SD. **f**). Western blot analysis of dystrophin, Lamin AC, and α-sarcoglycan protein expression in Tibialis Anterior 4 weeks post implantation, and histogram illustrating the level of dystrophin expression in TA muscle treated with U7-Mabs loaded scaffold in compare with normal TA muscle of wild type mouse. **Laminin = Lam, Lamin AC = LamAC, DYS = dystrophin.**

To quantify if U7-hMabs engraftment was maintained or changed overtime we measured the number of human Lamin AC^+^ cells in the back muscle, diaphragm and TA at four and eight weeks. Four weeks post-implantation, the number of human Lamin AC^+^ cells in back muscle was ∼ 30 cells/0.2mm^2^, which increased significantly at eight weeks ∼ 50 cells/0.2mm^2^ (p < 0.0001). In the diaphragm, a similar trend was observed, where the number of human Lamin AC^+^ cells increased from around 30 cells/0.2mm^2^ at four weeks to around 45 cells/0.2mm^2^ (p < 0.01) at eight weeks. Also, gastrocnemius muscle showed similar pattern of slight significant increase at 8 weeks (∼62 cells/0.2mm^2^) comparing with 4 weeks (∼57 cells/0.2mm^2^) (Supp Figure 6f). While, in the TA, and Tri the initial number of human Lamin AC^+^ cells (∼ 40, 67/0.2mm^2^ respectively) observed at four weeks post-implantation, remained stable at eight weeks, with no significant difference between time points (Figure 5d, Supp Figure 6f). In general, this progressive increase in donor cell numbers is likely, at least in part, attributable to the previously reported ability of U7-hMabs to improve the dystrophic muscle microenvironment, sufficient to guarantee survival of colonised muscle fibres, at variance with Wt-hMabs that did not restore dystrophin expression at a level sufficient for long term survival.

WB analysis of TA at four weeks post-implantation revealed robust dystrophin expression, reaching approximately 40% of healthy muscle levels, together with α-sarcoglycan expression, indicating effective reconstitution of the dystrophin-associated protein complex in mice receiving U7-hMabs (Figure 5d).

## Discussion

Using PCL nanofibers, we developed a 2D scaffold that supports cell adhesion and proliferation *in vitro* and *in vivo*, after sub-cutaneous implantation. In this study, we demonstrate that a Laminin-coated PCL (LamPCL) scaffold allows widespread delivery and engraftment of both Wt and U7-hMabs in many body muscles, resulting in significant dystrophin expression and reconstitution of the associated protein complex. This is a crucial first step towards easy, minimally invasive and affordable cell delivery for DMD but also for other cell therapies aiming at treating diseases affecting abundant and widespread tissues of our body. Considering the small size of the scaffold compared with the total mass of skeletal muscles, achieving a therapeutically significant and long-lasting engraftment remains a major challenge. However, the use of U7-hMabs enables a production of dystrophin of about 10% of a healthy muscle, not too lower than the presumed therapeutic threshold (19). This level of correction indicates that combining LamPCL scaffold-mediated delivery with exon-skipping–competent donor cells may at least in part overcome a main limitation of current cell and gene therapy approaches.

Beyond enabling localized cell delivery, a key feature of the LamPCL scaffold is its strong pro-angiogenic effect, which occurs independently of the presence of associated cells. The newly formed vascular network provide a gateway for donor cells to enter the systemic circulation, enabling dissemination to skeletal muscles distant from the implantation site, including TA, G and Tri, as well as muscles that are poorly targeted by conventional delivery methods, such as the diaphragm and heart.

Several mechanistic factors may contribute to this process: 1. Endothelial cells at the tip of the growing capillary lack a complete junctional apparatus facilitating entry of cells inside the lumen (20, 21); 2. Once the cells enter the capillary they are expected to follow the circulatory flow and enter the venous circulation with the risk of being trapped in the capillary filters such as the lung (22, 23). The detection of hMabs downstream of the capillary filters, where only few cells remain, indicate their ability to largely escape trapping. Supporting this interpretation, recent work has shown that cancer cells can flow through capillary filters as single cells while they are trapped when in clusters, thus giving rise to metastasis (24). In contrast to conventional intravascular delivery, where cells are administered at high concentrations and prone to aggregation, cells released gradually from the scaffold are likely to enter the circulation individually, facilitating transit through capillary filters.

This ability to mobilize cells into the circulation is particularly relevant for DMD, where targeting multiple muscle groups, including the diaphragm, is essential for therapeutic benefit.

Several challenges remain to be addressed before clinical translation. These include the durability of the scaffold, the exhaustion of the cell population, the possibility of repeated administrations and the need to efficiently treat the heart. The current scaffold formulation is resorbed after several months and although repeated minor surgical implantations are feasible, they are not ideal. We are currently optimizing the structure of the scaffold to extend its half-life, while the integration of a micro-branched port-a-cath system (PMID: 40540555) may enable repeated cell loading through a simple injection. Notably, repeated administration of hMabs is unlikely to pose immunological problems (9, 12). Efficient cardiac targeting is also essential, as improved skeletal muscle function may exacerbate underlying dilated cardiomyopathy if the heart is not concurrently treated. To this end we are developing a complementary approach using cardiac mesangioblasts (that do not undergo spontaneous cardiogenesis:(25), previously transduced with inducible cardiac transcription factors to promote cardiogenesis in situ.

Together, these advances lay the foundation for a transformative delivery platform that could eliminate the need for invasive catheter-based systemic administration, not only for DMD but also for other widespread monogenic diseases of the solid mesoderm, including most muscular dystrophies.

## Material and methods

### Scaffold fabrication and customization for cell culture

Polycaprolactone (PCL) with average Mn 80.000 g/mol (No. 440744, Sigma Aldrich, UK) was dissolved in 1,1,1,3,3,3-hexafluoro-2-propanol (HFIP; Sigma-Aldrich, United Kingdom) for 24 hours using magnetic stirrer at room temperature to prepare a 10% w/v of PCL solution (26). The prepared PCL solution was supplied by high voltage 20kv and electrospun fibres were collected at 20cm distance with flow rate 1ml/hr, and collector speed was set to be 1000 rpm for aligned fibre spinning.

The electro-spun sheet was mounted onto scaffoldex cell crown inserts - 48-well (Sigma Aldrich). Scaffold sterilization was done as previously described (27). To enhance cell attachment, the scaffold was coated for 2 hours with Laminin, (mouse, 1.2mg/ml) which diluted with PBS to be used in concentration (5ug/ml) (Thermo-Fisher Scientific). Scaffold was subsequently pre-wetted for 30 minutes – up to 24 hours in the culture medium depend on each tested cell line (28).

#### Scanning electron microscope analysis

The morphological characteristics of electrospun nanofiber was observed by scanning electron microscope (SBF-SEM Quanta Gatan 3View, Core Facility, Michael Smith, University of Manchester). Cell attachment on the scaffold was also analysed by SEM; wild type human mesangioblasts (Wt-hMabs) cells were plated on scaffold for 24 hours (n=3). Imaging was done with an accelerating voltage of 20 kV. Fibre diameters were measured using ImageJ software, and quantitative data were analysed using GraphPad Prism software, and normal distribution was assessed.

#### Cell types and culture conditions

For this study we used primary Wt-hMabs isolated from adult human muscle samples (Rectus Abdominis) or dystrophic Mabs (U7-Mabs) from the biopsy of a DMD patient. Both primary human cells were obtained from University Hospital South Manchester (UHSM), Manchester, UK with ethical approval (REC 13/SC/0499).

Mabs cells were maintained in hypoxic conditions (5%), plated at concentrations between 4000-6000 cells/cm^2^ and cultured in Mega-cell medium (M5), supplemented with 5% foetal bovine serum (Invitrogen), 1% L-Glutamine (2mM), 1% Non-Essential Amino Acids, 1% Pen/Strep, 0.2% B-Mercaptoethanol (0.1mM), 5ug/mL human FGF (Peprotech). To correct the genetic mutation, exon skipping was performed as previously described (1).

#### Cell biocompatibility, and proliferation

Live/Dead cell imaging kit (ThermoFisher-R37601), was used to check Wt-hMabs viability on scaffold.

To assess Wt-hMabs proliferation on the scaffold, cells were cultured in growth medium (M5) for 2, 5, and 10 days (n=3). Afterward, the cells were fixed and stained with Ki67 (Abcam, polyclonal rabbit) at a 1:200 dilution, followed by conjugation with Alexa Fluor 546 (Invitrogen). Myosin heavy chain MF20 (MyHC) (Development Study Hybridoma Bank) staining was also performed at a 1:2 dilution, conjugated to Alexa Fluor 488 (Invitrogen). A control group of cells cultured on a tissue culture plate (TCP), subjected to the same sterilization and laminin coating procedures as the LamPCL scaffold group, was included for comparison. Counted cells was analysed by GraphPad Prism Software and data were presented as mean ± standard deviation.

### In *vivo* scaffold implantation

Wild type immunodeficient NSG mice (thereafter will be refereed as Wt-NSG), and DMD51/HLA-A2/F725 strain, (thereafter will be referred as DMD-NSG) were produced by Jackson Laboratory and for DMD-NSG to contain the human sequence harbouring a deletion in exon 51 of the dystrophin gene (n=6 for each group) as described in (1).

The use of animals in this study was authorized by project license (PP8863485 and PL PDB0CF0C2). Anaesthesia was induced by 5% isoflurane before surgical procedure and maintained with 2.5%. Mice back skin was shaved and disinfected, bio-scaffold was implanted in a small subcutaneous pocket created by a short longitudinal incision (∼ 1 cm). The LamPCL scaffold was implanted on the right and left sides then fixed with 2-3 stiches of absorbable suture material; the skin was then closed with absorbable suture material (Vicryl 6-0), and pain killer (Buprenorphine 0.05-.1mg/kg) was administered once after surgery or more if needed. When any signs of infection appeared, antibiotics (Baytril 10% 85mg/kg/day) for 5 days was administered in water.

Wt-hMabs and U7-hMabs (200,000 cells) were seeded on LamPCL scaffold and kept for 24 hours in growth medium, then next day the scaffold was implanted over the back muscle of experimental mouse and surgical follow-up were performed for 5 days, then weekly close observations. Samples were collected after 2 weeks to observe scaffold vascularization, while 4, and 8 weeks to examine cell engraftment and dystrophin restoration.

### Immunofluorescence analysis

Cells and slides were fixed with 4% paraformaldehyde in PBS for either 15 minutes at room temperature, then were incubated in PBS containing 0.2% Triton X-100 (Sigma) and 1% Bovin serum albumin (BSA) (Sigma) for 30 minutes at room temperature. To block nonspecific antibody binding, sections were incubated in a blocking buffer consisting of PBS with 1% BSA, 0.2% Triton X-100, and 10% Foetal bovine serum (FBS) (GIBCO) for 1 hour at room temperature. Following blocking AffiniPure Fab fragment donkey anti-mouse IgG (Jackson immunoResearch) was added of the samples to avoid any unspecific binding with mouse tissue when mouse specific antibodies were used. Subsequently, primary antibody as anti-Ki67 (Ab 15580-Abcam), anti-MyHC MF20 (Development Studies, Hybridoma Bank), anti-F-actin (A12379-Invitrogen), anti-PECAM (ab182981 – Abcam), anti-Lamin AC (Mab636-Invitrogen), anti-Laminin (L9393-Sigma), anti-dystrophin (MANDRA17 (Development Studies Hybridoma Bank) incubated overnight in the blocking buffer. Secondary antibodies used included Alexa Fluor 488 anti-rabbit and Alexa Fluor 546 anti-mouse, and counter staining by DAPI.

The 400x magnification (40x oil objective, ∼0.2mm^2^) was selected for quantitative presentation and statistical analysis, as it provided clearer and more intense visualization of Lamin AC, and dystrophin expression compared to lower magnifications. IF images were captured by Confocal Lecia SP8, Bioimaging Facility, Michael Smith, University of Manchester).

### Western blot analysis

Tissues were carefully harvested and dissected into small pieces over dry ice before being transferred into round-bottom microcentrifuge tubes. Following this, each sample was snap-frozen in liquid nitrogen and stored at −80°C until required. To prepare the samples for analysis, a RIPA lysis buffer (10 mM TRIS, 100 mM NaCl, 1 mM EDTA, 1% Triton, 10% Glycerol, 0.1% SDS and 1% protease/phosphatase inhibitor) was added, and each sample was manually ground while submerged in liquid nitrogen to ensure complete homogenization. To determine protein concentration in the lysate, the Bradford assay (Bio-Rad protein assay) was employed. A 3-8% Tris-acetate gel was used for the electrophoresis, then the membranes were incubated with the following antibodies: anti-Lamin AC (MA3-1000 Invitrogen), anti-Dys MANDRA17 (Development Studies Hybridoma Bank), anti-Sarcoglycan (A007537) and anti-GAPDH (Ab 125247 Abcam). To visualize the protein bands, a chemiluminescent substrate (SuperSignal West femto substrate from Thermo Scientific) was applied to the membrane for several minutes. Chemiluminescent detection was then performed, capturing images of the protein bands and edited by using ImageLab software.

### Statistical analysis

All experiments in this project were performed at least in triplicate (biological), with three technical replicates in each. For microscopic imaging, a minimum of 20 images were captured per experiment. For quantification of Lamin AC and dystrophin, five images per sample were randomly selected for analysis. ImageJ software was used for cell counting, while GraphPad Prism software was utilized for statistical analysis. A two-tailed unpaired Student’s t-test with Welch’s correction was applied for comparisons between two groups, whereas one-way ANOVA was used for comparisons involving more than two groups, followed by Tukey’s or Sidak’s multiple comparison tests. Statistical significance is denoted by asterisks, with increasing numbers of asterisks indicating higher levels of significance (*p < 0.05), error bars = SD; data are presented as mean ± SD.

For the in *vivo* animal study, we adhered to the 3Rs principle (Replacement, Reduction, Refinement) to minimize animal use. A sample size of six animals (n= 6) was chosen to achieve statistical significance while ensuring the ethical reduction of animal use without compromising the study’s validity.

## Ethical statements

Anonymized adult human muscle samples (Rectus Abdominis) were obtained from University Hospital South Manchester (UHSM), Manchester, UK. All tissue samples were collected with informed consent from patients, and the procedures were approved by the UK National Research Ethics Committee (NRES), under protocol reference 13/SC/0499 (MA1, October 2015).

All in *vivo* experiments were conducted in accordance with the UK Animals (Scientific Procedures) Act 1986 (ASPA) and were approved by the Animal Welfare and Ethical Review Body (AWERB) and the Animals in Science Regulation Unit (ASRU). Experimental work was carried out under approved personal licence (PIL: 12172), project licences (PPL: PDB0CF0C and PP8863485) at the University of Manchester.

## Acknowledgements.

This work was supported by MRC grant R132096, the Wellcome Trust HICF, the ERC-AIG Unimab, the ERC PoC Unicardiomab to GC, the Astra Zeneca- MRC grant to FG, and by full scholarship [MM38/19] awarded to Amer. S. from The Egyptian Ministry of Higher Education & Scientific Research represented by The Egyptian Bureau for Cultural & Educational Affairs in London. The authors would like to thank Sarah Cartmell, Mohamed Elsawy, and the Henry Royce Institute for Advanced Materials (grant no. EP/R00661X/1, EP/S019367/1, EP/ P025021/1, and EP/P025498/1) for support and equipment use. We acknowledge the bioimaging, and electron microscopy (EM-FMBH) facilities at the University of Manchester for providing access to Confocal Leica SP8, and SEM which were instrumental in the completion of this research.

---

**Figure. Supp. 1:**
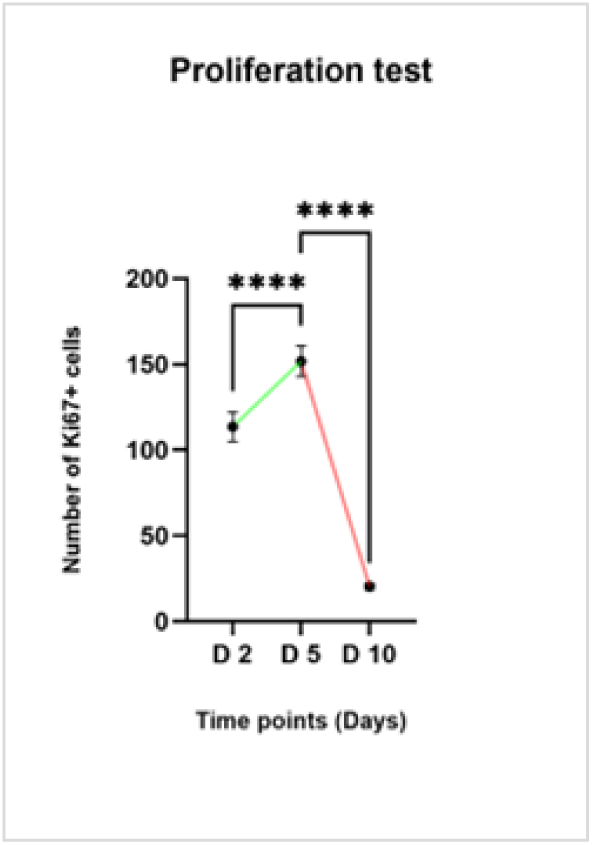
(a). Proliferation analysis of Wt-hMabs cultured on PCL nanofiber scaffolds, was assessed by counting of Ki67+ cells. The y-axis represents the number of Ki67+ cells, while the x-axis indicates different time points (2days, 5days, and 10days), n=3. Statistical analysis was performed using Ordinary One-Way ANOVA followed by Tukey’s multiple comparisons test. Data are presented as mean ± SEM. ****p < 0.0001 compared to other time points, error bars = SD; data are presented as mean ± SD.

**Figure. Supp. 2:**
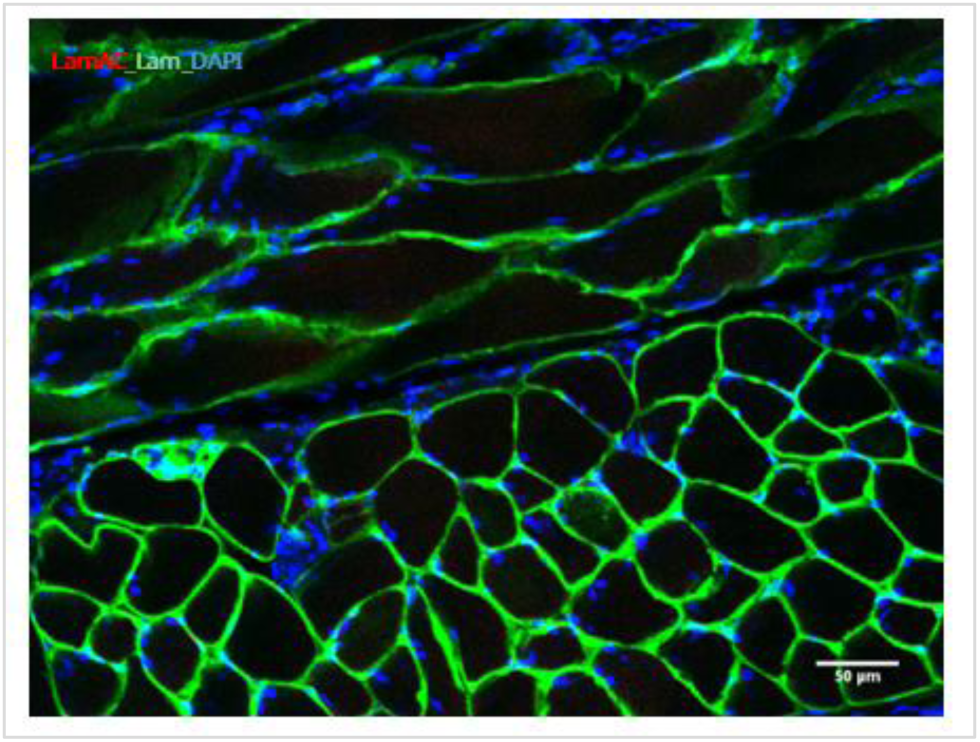
(a). Cell migration in WT-NSG mouse, 4 weeks post-implantation, representative immunofluorescence analysis of the underlying back muscle revealed absence of human Lamin AC+ cells, Lamin AC (red), Laminin (green), and DAPI (blue). **Laminin = Lam, Lamin AC = LamAC, DYS = dystrophin.**

**Figure. Supp. 3:**
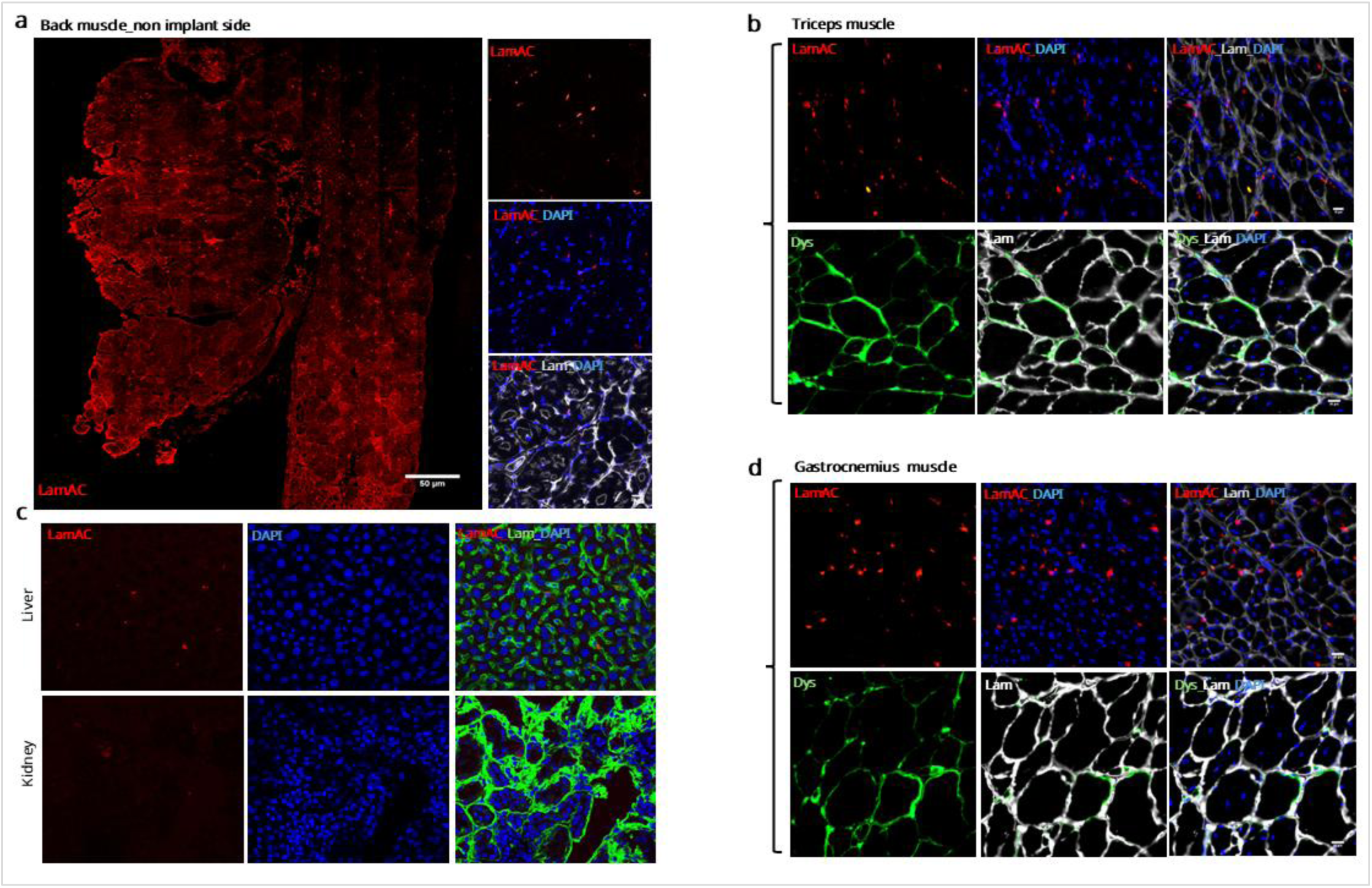
Engraftment and dystrophin restoration four weeks following transplantation of Wt-hMabs–loaded scaffolds in dystrophic mice. (**a**). Representative overview confocal microscopy images of a cross-sectioned non-implanted back muscle (contralateral side), showing distribution transplanted Wt-hMabs of back muscle-stained with antibodies against Lamin AC (red), Scale bar = 50µm. ***Vertical column,*** Higher magnification confocal images of back muscle stained with anti-Lamin AC (red), anti-laminin (grey), and DAPI (blue). (**b**). Representative confocal microscopy images of a cross-section of filter organs (liver, and kidney) showing the scattered presence of transplanted Wt-hMabs stained with anti-Lamin AC (red) laminin (green), and DAPI (blue). (**c**). Representative confocal microscopy images of a cross-section of triceps muscle (***upper row*)** showing the distribution of transplanted Wt-hMabs stained with anti-Lamin AC (red), anti-laminin (grey) antibodies, and DAPI (blue); **(*bottom row*)** shows dystrophin expression (green), Laminin (grey), and DAPI (blue), scale bar = 20µm. (**d**). Representative confocal microscopy images of a cross-sectioned of gastrocnemius muscle **(*upper row*)** showing the distribution of transplanted Wt-hMabs stained with anti-Lamin AC (red), anti-laminin (grey), and DAPI (blue), ***(bottom row),*** showing dystrophin expression (green), Laminin (grey), and DAPI (blue); scale bar = 20µm. **Laminin = Lam, Lamin AC = LamAC, DYS = dystrophin.**

**Figure. Supp. 4:**
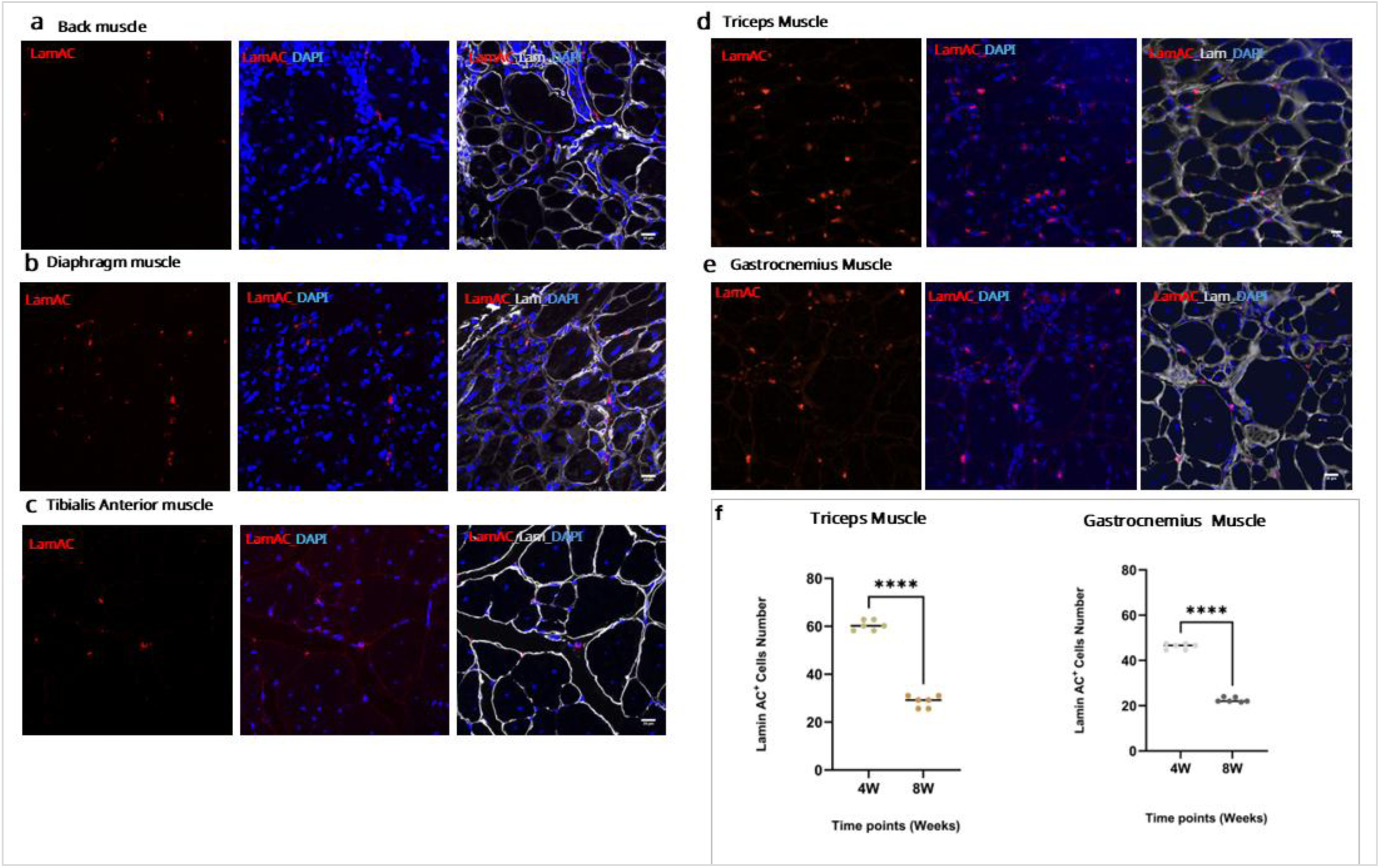
Wt-hMabs engraftment eight weeks following transplantation of Wt-hMabs loaded scaffolds in dystrophic mouse muscles. (**a**): back muscle; (**b**): diaphragm muscle; (**c**): Tibialis anterior muscle; (**d**): Triceps muscle; (**e**): Gastrocnemius muscle, all stained with anti-Lamin AC (red), anti-Laminin (grey) antibodies, and DAPI (blue); scale bar = 20µm. (**f**): Scatter blot graphs showing the number of Lamin AC+ cells distribution, in triceps, and gastrocnemius muscle at 4-, and 8weeks post-implantation with Wt-hMabs cells. The y-axis represents the number of Lamin AC+ cells/0.2mm^2^, while the x-axis indicates different time points (weeks). Each bar indicates the mean number of cells, n=6 represented by circles. Data analysis was performed using unpaired t-test with Welch’s correction, error bars = SD; data are presented as mean ± SD. Statistical significance is denoted by asterisks, with increasing numbers of asterisks indicating higher levels of significance (p < 0.05). **Laminin = Lam, Lamin AC = LamAC, DYS = dystrophin.**

**Figure. Supp. 5:**
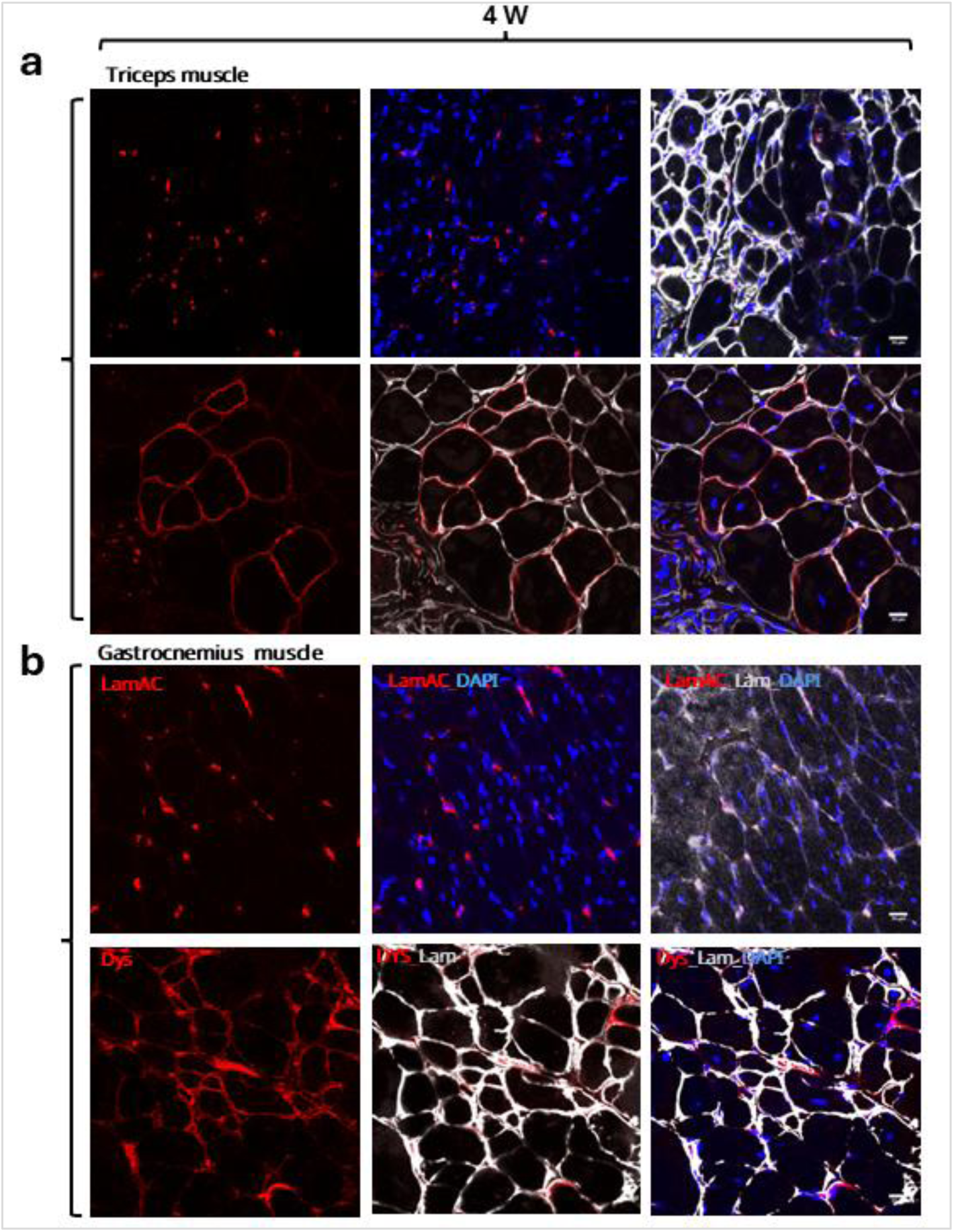
Engraftment and dystrophin restoration four weeks following transplantation of scaffolds loaded with dystrophic U7snRNA-genetically corrected human mesangioblast (U7-hMabs) in dystrophic mouse muscles. (**a**): Representative confocal microscopy images of a cross-section of triceps muscle, **upper row** shows the distribution of transplanted U7-hMabs stained with anti-Lamin AC (red), anti-laminin (grey) antibodies, and DAPI (blue); **bottom row** shows dystrophin expression (red), laminin (grey), and DAPI (blue), scale bar = 20μm. (**c**): Representative confocal microscopy image of a cross-section of gastrocnemius muscle, **upper row** shows the distribution of transplanted U7-hMabs stained with anti-Lamin AC (red), laminin (grey) antibodies, and DAPI (blue), **bottom raw** shows dystrophin expression (red), Laminin (grey), and DAPI (blue), scale bar = 20μm. **Laminin = Lam, Lamin AC = LamAC, DYS = dystrophin**

**Figure. Supp. 6:**
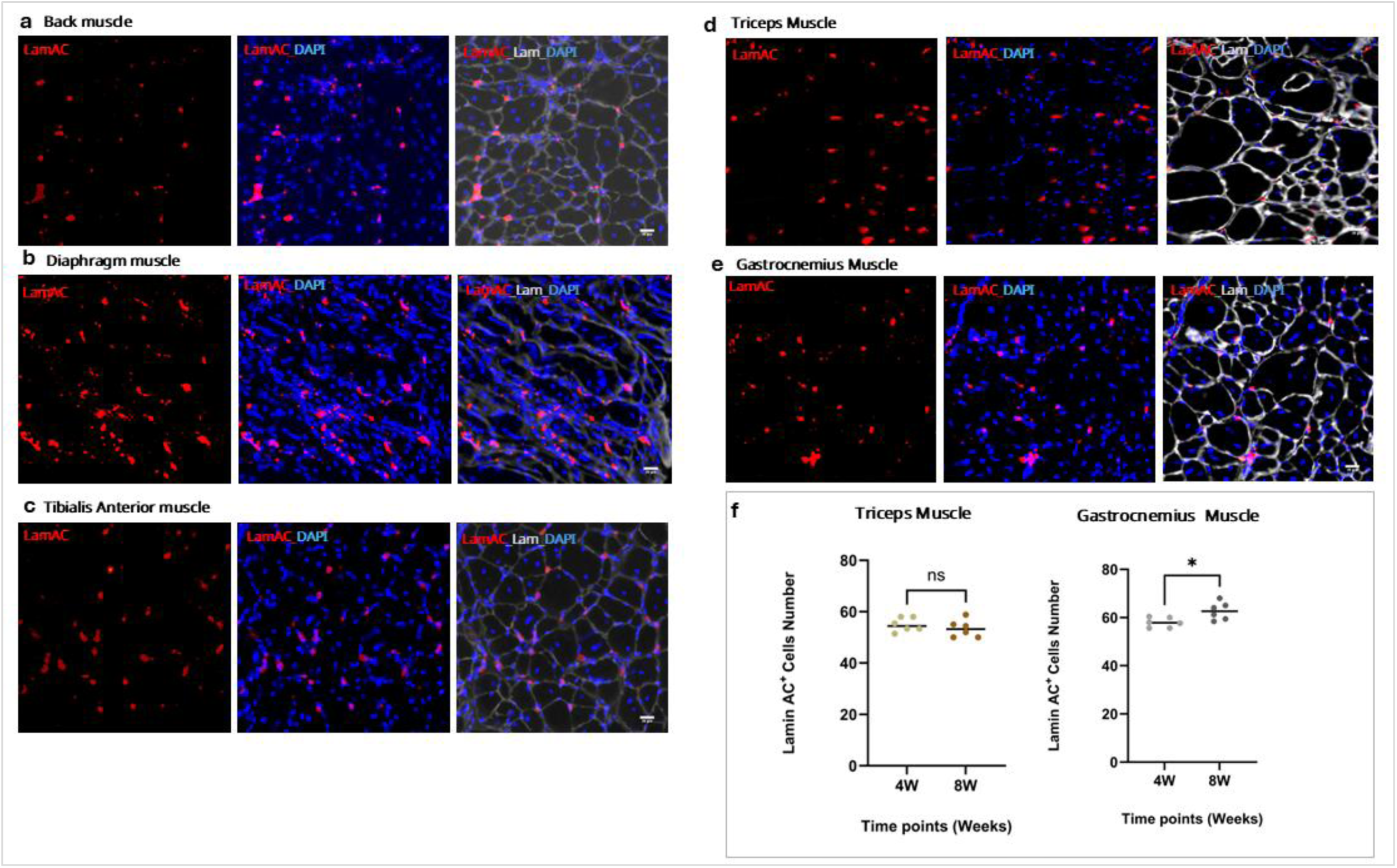
U7-hMabs engraftment eight weeks following transplantation of scaffolds loaded with dystrophic U7-hMabs cells in dystrophic mouse muscles, showing distribution of transplanted human cells in different muscle. (**a**): back muscle; (**b**): diaphragm muscle; (**c**): tibialis anterior muscle; (**d**): triceps muscle; (**e**): gastrocnemius muscle. All muscles were stained with anti-Lamin AC (red), anti-laminin (grey) antibodies and DAPI (blue); scale bar = 20µm. (**f**). Scatter blot graphs showing the number of Lamin AC+ cells distribution, in triceps, and gastrocnemius muscle at 4 and 8 weeks post-implantation with U7-hMabs. The y-axis represents the number of Lamin AC+ cells/0.2mm^2^, while the x-axis indicates different time points (weeks). Each bar indicates the mean number of cells, n=6 represented by circles. Data analysis was performed using unpaired t-test with Welch’s correction, error bars = SD; data are presented as mean ± SD. Statistical significance is denoted by asterisks, with increasing numbers of asterisks indicating higher levels of significance (p < 0.05). **Laminin = Lam, Lamin AC = LamAC, DYS = dystrophin.**

## Notes

### Competing Interest Statement

The authors have declared no competing interest.

## References.

1. Galli F, Bragg L, Rossi M, Proietti D, Perani L, Bacigaluppi M, et al. Cell-mediated exon skipping normalizes dystrophin expression and muscle function in a new mouse model of Duchenne Muscular Dystrophy. EMBO Molecular Medicine. 2024;16(4):927–44.

2. Koenig M, Hoffman EP, Bertelson CJ, Monaco AP, Feener C, Kunkel LM. Complete cloning of the duchenne muscular dystrophy (DMD) cDNA and preliminary genomic organization of the DMD gene in normal and affected individuals. Cell. 1987;50(3):509–17.

3. Tang A, Yokota T. Duchenne muscular dystrophy: promising early-stage clinical trials to watch. Expert Opinion on Investigational Drugs. 2024;33(3):201–17.

4. Oskoui M, Caller TA, Parsons JA, Servais L, Butterfield RJ, Bharadwaj J, et al. Delandistrogene moxeparvovec gene therapy in individuals with Duchenne muscular dystrophy: Evidence in focus: report of the AAN guidelines subcommittee. Neurology. 2025;104(11):e213604.

5. Philippidis A. Pfizer weighs next steps after DMD therapy linked to boy’s death fails phase III trial. Human Gene Therapy. 2024;35(13-14):413–5.

6. Duan D. Lethal immunotoxicity in high-dose systemic AAV therapy. Molecular Therapy. 2023;31(11):3123–6.

7. Akat A, Karaöz E. Cell Therapy Strategies on Duchenne Muscular Dystrophy: A Systematic Review of Clinical Applications. Stem Cell Reviews and Reports. 2024;20(1):138–58.

8. Sampaolesi M, Torrente Y, Innocenzi A, Tonlorenzi R, D’Antona G, Pellegrino MA, et al. Cell therapy of α-sarcoglycan null dystrophic mice through intra-arterial delivery of mesoangioblasts. Science. 2003;301(5632):487–92.

9. Sampaolesi M, Blot S, D’Antona G, Granger N, Tonlorenzi R, Innocenzi A, et al. Mesoangioblast stem cells ameliorate muscle function in dystrophic dogs. Nature. 2006;444(7119):574–9.

10. Tedesco FS, Hoshiya H, D’Antona G, Gerli MF, Messina G, Antonini S, et al. Stem cell–mediated transfer of a human artificial chromosome ameliorates muscular dystrophy. Science translational medicine. 2011;3(96):96ra78–96ra78.

11. Tedesco FS, Gerli MF, Perani L, Benedetti S, Ungaro F, Cassano M, et al. Transplantation of genetically corrected human iPSC-derived progenitors in mice with limb-girdle muscular dystrophy. Science translational medicine. 2012;4(140):140ra89–ra89.

12. Cossu G, Previtali SC, Napolitano S, Cicalese MP, Tedesco FS, Nicastro F, et al. Intra-arterial transplantation of HLA-matched donor mesoangioblasts in Duchenne muscular dystrophy. EMBO molecular medicine. 2015;7(12):1513–28.

13. Cossu G, Tonlorenzi R, Brunelli S, Sampaolesi M, Messina G, Azzoni E, et al. Mesoangioblasts at 20: From the embryonic aorta to the patient bed. Frontiers in genetics. 2023;13:1056114.

14. Schults JA, Young ER, Marsh N, Larsen E, Corley A, Ware RS, et al. Risk factors for arterial catheter failure and complications during critical care hospitalisation: a secondary analysis of a multisite, randomised trial. Journal of Intensive Care. 2024;12(1):12.

15. Perez-Puyana V, Wieringa P, Yuste Y, de la Portilla F, Guererro A, Romero A, Moroni L. Fabrication of hybrid scaffolds obtained from combinations of PCL with gelatin or collagen via electrospinning for skeletal muscle tissue engineering. Journal of Biomedical Materials Research Part A. 2021;109(9):1600–12.

16. Shi X, Cai A, Arkudas A, Horch RE, Jabeen S, Schubert DW, et al. Myoblast and ADSC coculture on conductive highly aligned nanofiber scaffolds for human skeletal muscle tissue engineering. Biomedical Materials. 2025;21(1):015003.

17. Galvez BG, Sampaolesi M, Brunelli S, Covarello D, Gavina M, Rossi B, et al. Complete repair of dystrophic skeletal muscle by mesoangioblasts with enhanced migration ability. The Journal of cell biology. 2006;174(2):231–43.

18. Galli D, Innocenzi A, Staszewsky L, Zanetta L, Sampaolesi M, Bai A, et al. Mesoangioblasts, Vessel-Associated Multipotent Stem Cells, Repair the Infarcted Heart by Multiple Cellular Mechanisms. Arteriosclerosis, Thrombosis, and Vascular Biology. 2005;25(4):692–7.

19. Godfrey C, Muses S, McClorey G, Wells KE, Coursindel T, Terry RL, et al. How much dystrophin is enough: the physiological consequences of different levels of dystrophin in the mdx mouse. Human molecular genetics. 2015;24(15):4225–37.

20. Eelen G, de Zeeuw P, Treps L, Harjes U, Wong BW, Carmeliet P. Endothelial cell metabolism. Physiological reviews. 2018;98(1):3–58.

21. Vincent Lc, Rafii S. Vascular frontiers without borders: multifaceted roles of platelet-derived growth factor (PDGF) in supporting postnatal angiogenesis and lymphangiogenesis. Cancer cell. 2004;6(4):307–9.

22. Fischer UM, Harting MT, Jimenez F, Monzon-Posadas WO, Xue H, Savitz SI, et al. Pulmonary passage is a major obstacle for intravenous stem cell delivery: the pulmonary first-pass effect. Stem Cells Dev. 2009;18(5):683–92.

23. Schrepfer S, Deuse T, Reichenspurner H, Fischbein MP, Robbins RC, Pelletier MP. Stem cell transplantation: the lung barrier. Transplant Proc. 2007;39(2):573–6.

24. Ring A, Nguyen-Straeuli BD, Wicki A, Aceto N. Biology, vulnerabilities and clinical applications of circulating tumour cells. Nature Reviews Cancer. 2023;23(2):95–111.

25. Galli D, Innocenzi A, Staszewsky L, Zanetta L, Sampaolesi M, Bai A, et al. Mesoangioblasts, vessel-associated multipotent stem cells, repair the infarcted heart by multiple cellular mechanisms: a comparison with bone marrow progenitors, fibroblasts, and endothelial cells. Arteriosclerosis, Thrombosis, and Vascular Biology. 2005;25(4):692–7.

26. Shamsah AH, Cartmell SH, Richardson SM, Bosworth LA. Material characterization of PCL: PLLA electrospun fibers following six months degradation in vitro. Polymers. 2020;12(3):700.

27. Bhaskar P, Bosworth LA, Wong R, O’brien MA, Kriel H, Smit E, et al. Cell response to sterilized electrospun poly (ɛ-caprolactone) scaffolds to aid tendon regeneration in vivo. Journal of Biomedical Materials Research Part A. 2017;105(2):389–97.

28. Phamornnak C, Han B, Spencer BF, Ashton MD, Blanford CF, Hardy JG, et al. Instructive electroactive electrospun silk fibroin-based biomaterials for peripheral nerve tissue engineering. Biomaterials Advances. 2022;141:213094.

